# AlphaFast: High-throughput AlphaFold 3 via GPU-accelerated MSA construction

**DOI:** 10.64898/2026.02.17.706409

**Authors:** Benjamin C. Perry, Jeonghyeon Kim, Philip A. Romero

**Author notes:** These authors contributed equally to this work (Benjamin C. Perry, Jeonghyeon Kim). Contributing authors.

## Abstract

AlphaFold 3 (AF3) enables accurate biomolecular modeling but is limited by slow, CPU-bound multiple sequence alignment (MSA) generation. We introduce AlphaFast, a drop-in framework that integrates GPU-accelerated MMseqs2 sequence search to remove this bottleneck. AlphaFast achieves a 68.5× speedup in MSA construction and a 22.8× reduction in end-to-end runtime on a single GPU, and delivers predictions in 8 seconds per input on four GPUs while maintaining indistinguishable structural accuracy. A serverless deployment enables structure prediction for as little as $0.035 per input. Code is available at https://github.com/RomeroLab/alphafast

---

The release of AlphaFold 3 (AF3) extended accurate protein modeling from protein chains to protein–ligand, protein–DNA, and protein–RNA complexes [1]. Despite its predictive power, inference remains computationally expensive, limiting practical use in high-throughput experiments in proteomics, interactomics, or synthetic biodesign. A key contributor to this cost is the construction of multiple sequence alignments (MSAs). MSAs encode evolutionary information that is critical to state-of-the-art folding and interaction prediction, but their generation requires searching giga-scale reference databases [2].

Over the past few decades, MSA construction has progressed from exact dynamic programming solutions to fast heuristic and sensitive profile-based aligners such as BLAST [3], DIAMOND [4], HMMER [5], and MMSeqs [6, 7]. While these advances substantially reduce search time, large-scale MSA generation remains computationally demanding. As a result, many modern biomolecular prediction pipelines that strictly require [8–10] or obtain best results from inclusion of MSAs [11] outsource their construction to hosted web servers [10, 12]. While this approach lowers the barrier to entry, it introduces additional latency, prevents custom MSA parameter or database tuning, complicates deployment in high-performance computing (HPC) environments, and remains low throughput.

Recently, MMseqs2-GPU demonstrated that GPU-accelerated sequence search can achieve an order-of-magnitude speedup over CPU-bound pipelines, enabling fast, fully local MSA generation while substantially reducing inference time in AlphaFold 2 workflows [13]. However, extending these gains to AF3 is non-trivial given incompatible data pipelines, the post- and pre-processing requirements of MMSeqs2-GPU, as well as file input/output (I/O) optimization in HPC environments. Furthermore, there exists no cost-effective, high throughput implementation of AF3 available to researchers without access to significant computational resources.

In this work, we introduce AlphaFast, a simplified pipeline that replaces CPU-bound JackHMMER with GPU-accelerated MMseqs2 while preserving the original AF3 folding module, weights, and end-to-end performance. AlphaFast supports cached batch processing for ligands and proteins in both single-GPU and multi-GPU setups. On a single GPU, AlphaFast achieves up to a 68.5*×* speedup in MSA generation and a 22.8*×* reduction in end-to-end (wall clock). With four GPUs, AlphaFast reaches speeds of 8 seconds per input (a 71.2*×* acceleration), scaling nearly linearly with additional GPUs. We verify that AlphaFast produces MSAs and structures that are statistically indistinguishable from AF3 MSAs and outputs. Finally, we note this modular design is extensible and should serve as an implementation framework to any folding architecture that decouples MSA construction from inference.

AlphaFast introduces three key architectural improvements to AF3. First, unlike AF3’s per-chain JackHMMER search (Figure 1a, top), AlphaFast consolidates unique sequences into a batched MMseqs2-GPU query for sequential database search (Figure 1a, bottom). Second, AlphaFast maximizes throughput by offloading MSA post-processing of database *N* concurrently with GPU search of database *N* + 1. Finally, AlphaFast enforces a strict two-stage architecture to resolve video memory (VRAM) conflicts between JAX initialization [14] and MSA generation. For fair comparison, we strictly match AF3’s default E-values (10^*−*4^) and sequence limits for all four reference databases: UniRef90 [15], MGnify [16], Small BFD [8, 10], and UniProt [17].

**Fig. 1.**
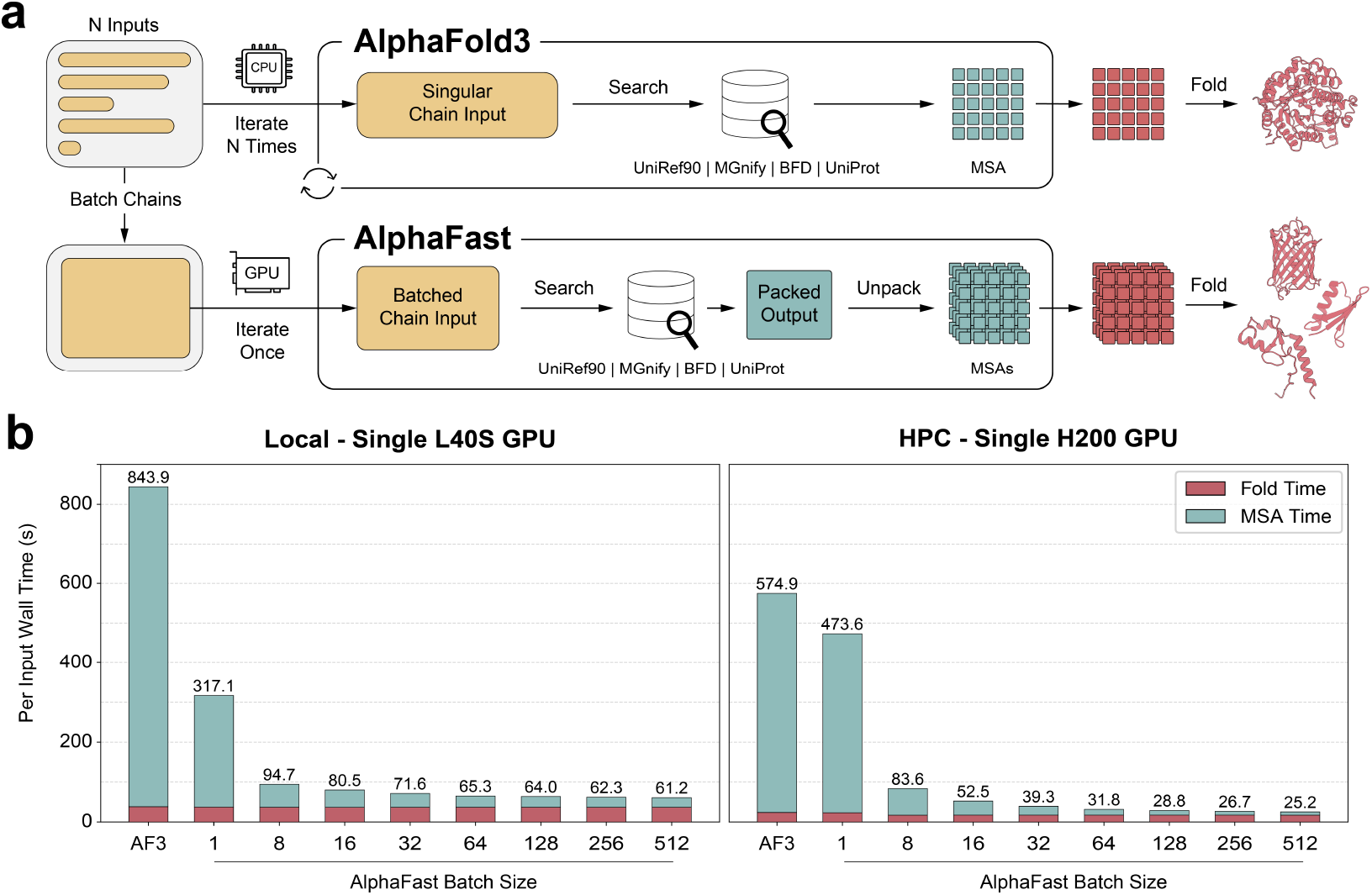
AlphaFast architecture enables inference speeds scaling with batch size. (a) Architectural comparison between AF3 (top) and AlphaFast (bottom). AF3 iterates *N* times over inputs on CPU, processing one chain at a time through parallel database searches. AlphaFast batches all chains, iterates once on GPU, searches sequentially, produces a packed output database, then unpacks and converts individual MSAs before folding. (b) Average total wall time running inference a single NVIDIA L40S GPU on a bare metal server with Docker (left) vs a single NVIDIA H200 GPU on an HPC environment with Singularity (right). Stacked bar charts demonstrate homology search dominates AF3 inference and is increasingly reduced at larger query batch sizes.

To benchmark AlphaFast against AF3, we curated a set of 32 and 512 protein monomer, protein-protein, and protein-ligand targets sampled from the PDB [18] (see the Supplementary Information for selection criteria and chain-length distributions). For each configuration, we ran benchmarks in triplicate, measuring average per-input wall time while varying the AlphaFast GPU query batch size from 1 to 512. For fair comparison, we run AF3 in batched mode with 4 parallel JackHMMER searches and model compilation. We test AlphaFast in two common setups: a local, bare metal server with NVIDIA L40S GPUs and an HPC environment with the SLURM job scheduler [19] and NVIDIA H200 GPUs. AlphaFast speedups are calculated relative to AF3 running on the exact same hardware.

On a single NVIDIA L40S GPU (Figure 1b, left), AF3 requires 843.9 seconds per input, with MSA construction accounting for the vast majority (95.4%) of wall time. Without batching, AlphaFast reduces this to 317.1 seconds and continues to improve with larger batch sizes. Efficiency gains begin to plateau around a batch size of 64 (65.3 seconds per input), likely due to GPU memory bandwidth saturation. The highest efficiency gain occurs at the maximum tested batch size of 512 (61.2 seconds per input), yielding a maximum 13.8*×* end-to-end speedup on a single L40S GPU. Larger batch sizes likely will reduce this further, but diminishing returns are expected.

On a single NVIDIA H200 GPU, performance gains are amplified further (Figure 1b, right). The increased HBM3e memory bandwidth enables continued scaling across all tested batch sizes without a clear plateau in MSA per-input time. AF3 requires 574.9 seconds per input, with MSA construction consuming 95.8% of total wall time. Wall time decreases rapidly to a minimum of 25.2 seconds per input at the maximal tested batch size of 512 (a 22.8*×* speedup). At this batch size, MSA time consumes 31.9% of runtime and is likely to reduce further at even larger batch sizes. Notably, sequential processing yields only a 1.2-fold speedup on the H200, slower than the L40S at the same batch size. We attribute this to I/O bottlenecks on the shared HPC filesystem during intermediate file production; at larger batch sizes, GPU throughput compensates for this overhead.

For multi-GPU inference, AlphaFast employs a phase-separated parallel architecture. In Phase 1, inputs are partitioned round-robin across all available GPUs and each GPU independently runs the batched MSA pipeline on its partition. After all MSA processes complete, intermediate feature files are written to disk. Phase 2 then re-partitions the outputs across GPUs for parallel folding. This design ensures clean separation of GPU-bound MSA search from JAX-based inference while achieving near-linear scaling. We expand the test set to 2, 048 inputs to account for a batch size of 512 on 4 GPUs (see Supplementary Information). On a 4xL40S setup, AlphaFast achieves a throughput of 19.4 seconds per input (8.2 second MSA, 11.2 second fold), a 3.1*×* improvement over the single-GPU configuration and a 43.5*×* end-to-end speedup over the AF3 baseline. On a 4xH200 setup, per-input time drops further to 8.1 seconds per input (3.3 seconds MSA, 4.8 second fold), yielding a 71.2*×* speedup over the same-hardware baseline and reducing MSA construction time by over two orders of magnitude. Both configurations exhibit approximately 78% parallel efficiency across four GPUs. The remaining overhead is likely attributable to phase synchronization and I/O. Because the two phases are independent and require no inter-GPU communication, throughput is expected to scale near-linearly with additional GPUs.

To democratize access to high-throughput structure prediction, AlphaFast includes a serverless inference mode via the compute provider Modal. In this mode, users only require a copy of the AlphaFast weights, a billing account, and a directory of inputs. We benchmarked performance on both NVIDIA L40S and H200 GPUs to optimize for cost-efficiency. Remarkably, despite the H200’s significantly higher hourly rental rate compared to the L40S, it proved to be the more economical choice due to its superior throughput. The H200 achieved a 2.3× speedup, reducing the inference time to 28.3 seconds and the total cost to $0.035 per target (lower than the L40S). Detailed cost benchmarks are provided in the Supplementary Information.

To evaluate the numerical performance of AlphaFast relative to AF3, we used a Two One-Sided T-test (TOST) [20, 21]. We applied distinct metrics and criteria to reflect the biological nature of each data type. For MSA inputs (*N*_eff_, Depth), we performed a Log-Ratio TOST against the standard bioequivalence margin of [0.80, 1.25]. For structural outputs (TM-score, RMSD), we assessed the absolute mean difference (Δ) against strict margins (e.g., *±* 0.02 for TM-score).

Input consistency is visualized in Figure 2a-b. While AlphaFast retrieves fewer raw sequences (MSA Depth GMR *≈* 87.1%), it successfully captures the full spectrum of effective evolutionary information (*N*_eff_ GMR *≈*107.6%). As detailed in the Supplementary Information, these values fall well within or exceed the bioequivalence thresholds, confirming that AlphaFast selectively filters redundant sequences without compromising the information density required for prediction.

**Fig. 2.**
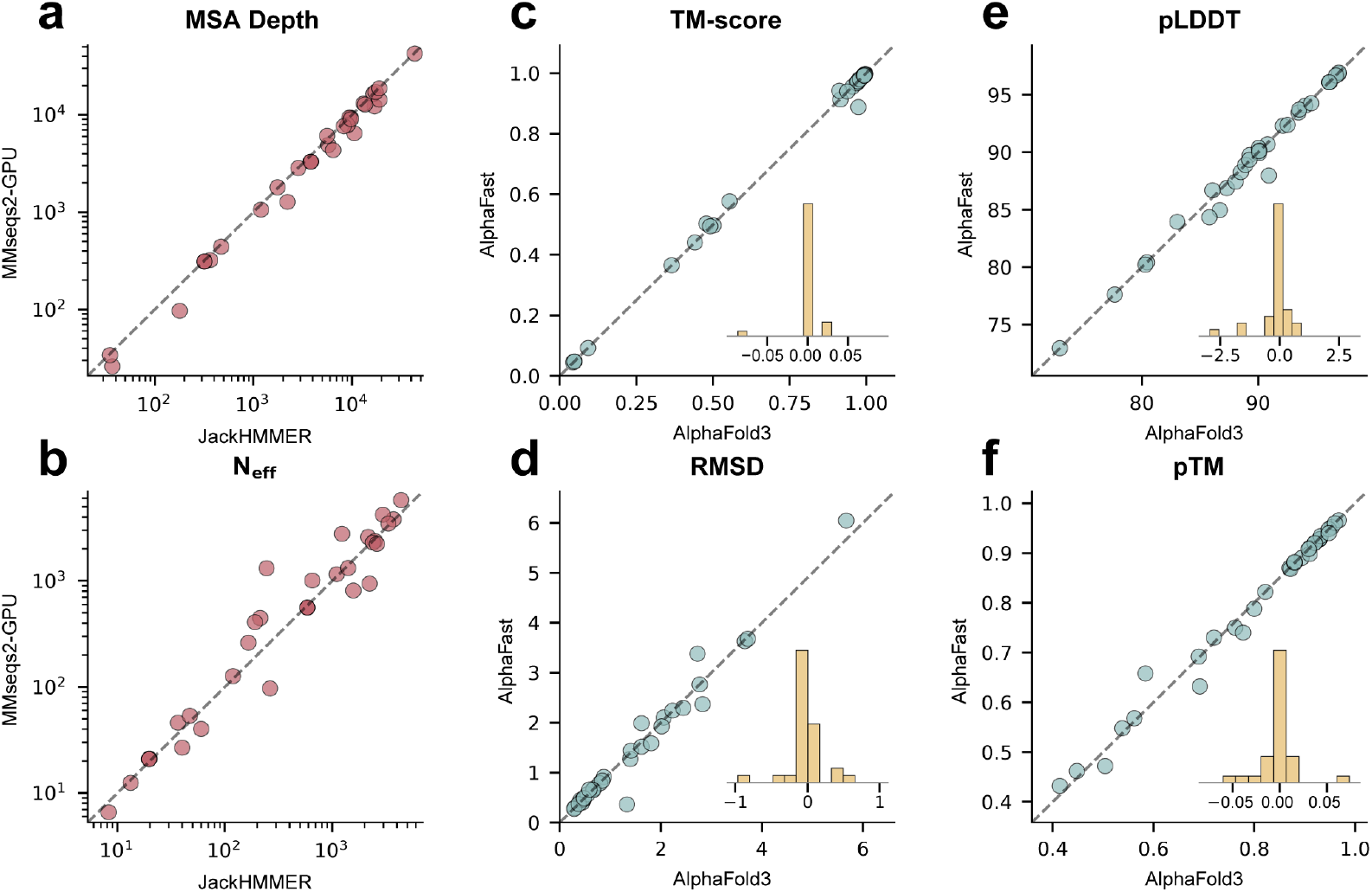
AlphaFast achieves structural prediction accuracy equivalent to AF3. (a-b) Log-log scatter plots of MSA Depth and *N*_eff_. (c-d) Global structural accuracy (TM-score and RMSD) shows high concordance (*y* = *x*). Inset histograms display the distribution of pairwise differences (Δ = AlphaFast*−*AF3), showing a sharp peak at zero. (e-f) Model confidence metrics (pLDDT and pTM) demonstrate equivalence within bounds. While MSA Depth is reduced, effective information content (*N*_eff_) and structural metrics remain statistically indistinguishable.

Consequently, this input efficiency translates directly to high-fidelity structural outputs. As shown in Figure 2c-f, the variance between methods is negligible. The mean difference for TM-score was virtually zero (Δ *≈* +0.002), and RMSD showed no deviation (Δ*≈* 0.00 Å). These results demonstrate that AlphaFast maintains strict structural equivalence to the baseline despite the optimization of input generation.

In conclusion, AlphaFast overcomes the computational bottleneck of CPU-bound MSA generation in AlphaFold 3 by integrating a GPU-accelerated search via MMseqs2. This approach decouples feature generation from inference, delivering a massive 22.8*×* speedup on a single GPU and up to 71.2*×* in multi-GPU configurations, effectively reducing per-target runtime from nearly 20 minutes to under 10 seconds. Crucially, this acceleration preserves the quality of evolutionary information (*N*_eff_) and structural accuracy, ensuring that speed does not come at the cost of model fidelity. To democratize access to this high-throughput capability, we provide a serverless implementation enabling researchers to fold large cohorts for approximately *∼* $0.035 per target.

Despite these advances, AlphaFast has specific limitations. Efficiency gains are maximized for large batches of unique proteins; workflows centered on a single static target—such as repetitive small molecule docking against one receptor—may not realize the same benefits due to lack of caching opportunities. Furthermore, performance on extreme sequence lengths or non-natural inputs remains to be fully characterized. Looking forward, the strategy of separating search from inference provides a generalizable template for removing bottlenecks in other structural biology models. Ultimately, AlphaFast demonstrates that using the right tools virtually eliminates MSA lookup time, enabling industrial-scale protein design for academic laboratories.

## Supplementary information

The online version contains supplementary material. This file includes detailed hardware specifications, full experimental scaling datasets, extended statistical bioequivalence assessments, and additional figures regarding dataset distribution and template retrieval performance.

## Acknowledgements

We acknowledge and thank the Duke Compute Cluster and Pratt Information Technology office for providing and managing computing resources that have contributed to the research results reported within this paper. Furthermore, we thank members of the Romero lab for helpful discussions and figure preparation assistance related to this work.

## Declarations

### Data availability

The benchmark datasets generated and analyzed during the current study are available in the Figshare repository, https://doi.org/10.6084/m9.figshare.31343287.

### Code availability

The source code for AlphaFast is openly available on GitHub at https://github.com/RomeroLab/alphafast.

### Author contribution

**Benjamin Perry:** Conceptualization, Methodology, Software, Investigation, Formal Analysis, Visualization, Writing – Original Draft. **Jeonghyeon Kim:** Methodology, Software, Investigation, Formal Analysis, Visualization, Writing – Original Draft. **Philip Romero:** Conceptualization, Writing – Review & Editing, Supervision, Project Administration, Funding Acquisition.

## References

[1] Abramson, J. et al. Accurate structure prediction of biomolecular interactions with alphafold 3. Nature 630, 493–500 (2024).

[2] Lee, S. et al. Petabase-scale homology search for structure prediction. Cold Spring Harbor perspectives in biology 16, a041465 (2024).

[3] Altschul, S. F., Gish, W., Miller, W., Myers, E. W. & Lipman, D. J. Basic local alignment search tool. J. Mol. Biol. 215, 403–410 (1990). URL 10.1016/S0022-2836(05)80360-2.

[4] Buchfink, B., Reuter, K. & Drost, H.-G. Sensitive protein alignments at tree-of-life scale using DIAMOND. Nat. Methods 18, 366–368 (2021). URL https://www.nature.com/articles/s41592-021-01101-x.

[5] Eddy, S. R. Accelerated profile hmm searches. PLoS computational biology 7, e1002195 (2011).

[6] Hauser, M., Steinegger, M. & Söding, J. MMseqs software suite for fast and deep clustering and searching of large protein sequence sets. Bioinformatics 32, 1323–1330 (2016). URL 10.1093/bioinformatics/btw006.

[7] Steinegger, M. & Söding, J. Mmseqs2 enables sensitive protein sequence searching for the analysis of massive data sets. Nature biotechnology 35, 1026–1028 (2017).

[8] Jumper, J. et al. Highly accurate protein structure prediction with alphafold. nature 596, 583–589 (2021).

[9] Wohlwend, J. et al. Boltz-1 democratizing biomolecular interaction modeling. bioRxivorg (2025). URL 10.1101/2024.11.19.624167.

[10] Mirdita, M. et al. ColabFold: making protein folding accessible to all. Nat. Methods 19, 679–682 (2022). URL https://www.nature.com/articles/s41592-022-01488-1.

[11] Discovery, C. et al. Chai-1: Decoding the molecular interactions of life. bioRxiv (2024). URL 10.1101/2024.10.10.615955.

[12] Mirdita, M., Steinegger, M. & Söding, J. MMseqs2 desktop and local web server app for fast, interactive sequence searches. Bioinformatics 35, 2856–2858 (2019). URL 10.1093/bioinformatics/bty1057.

[13] Kallenborn, F. et al. GPU-accelerated homology search with MMseqs2. Nat. Methods 22, 2024–2027 (2025). URL https://www.nature.com/articles/s41592-025-02819-8.

[14] Bradbury, J. et al. Jax: composable transformations of python+ numpy programs (2018).

[15] Suzek, B. E. et al. Uniref clusters: a comprehensive and scalable alternative for improving sequence similarity searches. Bioinformatics 31, 926–932 (2015).

[16] Mitchell, A. L. et al. Mgnify: the microbiome analysis resource in 2020. Nucleic acids research 48, D570–D578 (2020).

[17] Consortium, U. Uniprot: a worldwide hub of protein knowledge. Nucleic acids research 47, D506–D515 (2019).

[18] wwPDB consortium. Protein data bank: the single global archive for 3d macromolecular structure data. Nucleic Acids Research 47, D520–D528 (2018). URL 10.1093/nar/gky949.

[19] Yoo, A. B., Jette, M. A. & Grondona, M. SLURM: Simple linux utility for resource management (2003). URL 10.1007/109689873.

[20] Koch, G. G. One-sided and two-sided tests and ρ values. Journal of biopharmaceutical statistics 1, 161–170 (1991).

[21] Schuirmann, D. J. A comparison of the two one-sided tests procedure and the power approach for assessing the equivalence of average bioavailability. Journal of pharmacokinetics and biopharmaceutics 15, 657–680 (1987).

