## Supplementary Information for "AlphaFast: High-throughput AlphaFold 3 via GPU-accelerated MSA construction"

### Appendix A Benchmark Data

#### A.1 Reference Sequence Data & Preparation

For benchmarking, we acquire reference sequence databases identical to those fetched in AlphaFold 3 (AF3) via the `fetch_databases.sh` script. The raw sequence data were obtained in FASTA format, including UniRef90 v2022_05 [1], MGnify clusters v2022_05 [2], BFD first non-consensus sequences [3, 4], and UniProt v2021_04 [5]. These represent the four databases searched during MSA lookup in the AF3 pipeline. JackHMMER [6] can utilize the FASTA (.fa) format without any further pre-processing; However, MMSeqs2-GPU [7] requires databases to be converted into a memory-aligned "padded" binary format first. We implemented a two-step conversion process using MMseqs2: first converting FASTA files to binary, indexed databases, and subsequently generating padded sequence databases optimized for GPU memory access. We use the following commands per reference sequence database using 128 threads for speed:

```
# 1. Convert FASTA to standard MMseqs2 database
mmseqs createdb <database-name>.fa <database-name>

# 2. Generate Padded DB for GPU acceleration
mmseqs makepaddedseqdb <database-name> <database-name>_padded>
```

The resulting `*_padded` databases reduce the need for loading massive index files into RAM, instead streamlining data transfer to the GPU during inference. Below, we enumerate the disk space differences between database formats. For MMSeqs2-GPU databases, we sum the space used by the lookup, index, type, header and main database files.

**Table A1 Database Infrastructure Comparison: Raw FASTA vs. GPU-Padded Formats.** The table compares the storage footprint of the original raw sequences against the converted padded databases required for GPU acceleration. While the padded format increases storage usage by 27.3% due to memory alignment, it is strictly required.

| Database | Raw FASTA Size (GiB) | GPU Padded Size (GiB) | Role |
|---|---|---|---|
| UniRef90 | 66.9 | 78.8 | MSA Generation |
| MGnify | 119.7 | 169.9 | MSA Generation |
| Small BFD | 16.9 | 22.1 | MSA Generation |
| UniProt (Paired) | 101.0 | 116.7 | Paired MSA |
| **Total** | **304.5** | **387.5** | **$\Delta$ = 83 GiB** |

On our local system, the data setup took approximately 4 hours for download and conversion. However, we note actual times may vary quite significantly depending on several factors including but not limited to: download speed, system write input/output (I/O), host server API call limits, and download agent (`wget` vs. `aria2c`).

#### A.2 Scaling & Numerical Performance Dataset Preparation

To ensure rigorous evaluation of both the standard and accelerated pipelines, we curated benchmark datasets from the Protein Data Bank [8]. We selected structures released after September 30, 2021, aligning with the training data cutoff of AF3 to prevent data leakage. The curation process excluded DNA/RNA complexes to focus strictly on protein targets. We applied the following filtering criteria to ensure high-quality structural data:

1. **Resolution:** Structures with resolution $\leq 3.0$ Å (including NMR/EM structures).
2. **Chain Length:** Individual chains between 50 and 500 residues.
3. **Total Size:** Total residue count $\leq 1,500$ to fit within standard GPU memory constraints.
4. **Complex Types:** Targets were categorized into three groups: monomers (single chain), protein-ligand (single chain with small molecules), and protein-protein complexes (multimers).

#### A.3 Multi-Tiered Dataset Strategy

Given the substantial computational resources required for end-to-end AF3 inference, we adopted a multi-tiered dataset strategy. This approach allows us to efficiently evaluate prediction accuracy, single-GPU throughput, and multi-GPU scalability without incurring prohibitive computational costs. All datasets were randomly sampled from the filtered PDB criteria to ensure they share representative structural characteristics (e.g., chain length, complex type distribution) without systematic bias. The exception is the multi-GPU benchmark set, which we relax the training cutoff date.

1. **Accuracy Benchmark ($N = 32$).** For the primary evaluation of MSA quality and structure prediction accuracy (Section B), we utilized a subset of 32 randomly sampled targets. The size of this dataset was constrained by the extreme computational cost of the baseline pipeline; generating reference MSAs and structures using the standard AF3 (JackHMMER) pipeline requires approximately 20 minutes per target on our hardware. A larger accuracy benchmark would have necessitated prohibitively long runtimes solely to establish the baseline ground truth.
2. **Single-GPU Benchmark ($N = 512$).** To rigorously evaluate throughput, latency, and cost-efficiency on single-device configurations (e.g., NVIDIA L40S vs. H200), we curated a mid-sized dataset of 512 targets. This volume is sufficient to saturate the GPU and mitigate cold-start overheads, providing a robust estimate for "production-grade" performance and serverless cost calculations (Section C.3).
3. **Multi-GPU Benchmark ($N = 2,048$).** For the most demanding stress tests, including multi-GPU scalability and storage I/O bottlenecks in an High Performance Computing (HPC) environment, we constructed a large-scale dataset of 2,048 targets. This tier validates the stability of the multi-gpu architecture under heavy load and demonstrates the system's capability to handle industrial-scale screening workflows. For this set only, we relax the training data cutoff to structures released after September 30, 2016 as our sampling strategy did not yield enough inputs to test this scale. We only utilize this large-scale set to assess inference speed and not numerical performance.

### Appendix B Architecture & Evaluation Methodology

#### B.1 Homology and Template Search

To benchmark the performance of the accelerated pipeline, we established a controlled comparison between the default AF3 data generation process and our proposed MMseqs2-GPU implementation. Both pipelines search against identical sequence databases with attempts to match parameters to ensure comparable MSA composition.

1. **Baseline Pipeline (JackHMMER/hmmsearch).** The default pipeline utilized HMMER 3.4 for MSA generation. Four parallel subprocess calls query against sequence databases with the following per-database sequence limits:
   (a) **UniRef90.** Maximum of 10,000 sequences
   (b) **MGnify.** Maximum of 5,000 sequences
   (c) **Small BFD.** Maximum of 5,000 sequences
   (d) **UniProt.** Maximum 50,000 sequences (for taxonomic pairing)

   Search parameters were configured with a single iteration ($N = 1$), E-value threshold of $10^{-4}$, and aggressive pre-filters ($F1 = 5 \times 10^{-4}$, $F2 = 5 \times 10^{-5}$, $F3 = 5 \times 10^{-7}$) to accelerate queries against large metagenomic databases. For template retrieval, HMM profiles constructed from the merged MSA were searched against the `pdb_seqres` database using `hmmsearch`, configured with relaxed pre-filters ($F1 = F2 = F3 = 0.1$) and an E-value threshold of $10^{-3}$.
2. **Accelerated Pipeline (MMseqs2-GPU).** The experimental pipeline utilized MMSeqs2 with GPU-accelerated pre-filtering (`--gpu 1`) at sensitivity $s = 7.5$. To maximize GPU utilization while avoiding memory contention, we implemented a pipelined execution strategy:
   (a) **Sequential GPU Search.** Database searches execute sequentially against padded databases (UniRef90, MGnify, Small BFD, UniProt), with each `mmseqs search` command using alignment backtraces (`-a`) for downstream MSA generation. Sequence limits match the baseline pipeline with $E \leq 10^{-4}$.
   (b) **Parallel CPU Post-Processing.** Upon completion of each GPU search, post-processing tasks (`mmseqs result2msa` with `--msa-format-mode 5` for A3M output, followed by `mmseqs unpackdb`) are offloaded to a thread pool executor, allowing CPU-bound conversion to overlap with subsequent GPU searches.

(c) **MSA Consolidation.** Individual database MSAs are merged with feature-equivalence deduplication to remove redundant sequences across databases.

For template retrieval, `mmseqs search` queries the `pdb_seqres` database ($E \leq 10^{-3}$), filtering for minimum alignment coverage of 40%. Template metadata (PDB ID, chain, residue ranges) are parsed from result alignments to map hits to mmCIF structures, excluding entries released after September 30, 2021 for fair comparison.

##### B.2 Folding Method & Template Filtering

Structure prediction was performed using the official **JAX-based** [9] implementation of AF3 within a custom Docker environment (Singularity was used on the HPC environment instead). To capture the model's stochasticity and robustly estimate performance, we generated a total of 25 predictions per target, derived from 5 distinct random seeds with 5 diffusion samples per seed. For scaling, we reduce this to 5 predictions per target, derived from 1 random seed with 5 diffusion samples. Inference was executed with standard hyperparameters, including 10 recycling iterations and Triton-based flash attention [10].

To evaluate performance under controlled evolutionary contexts, we implemented two distinct template regimes by pre-filtering the template inputs before inference:

1. **Relaxed Regime (ID $<$ 90%).** A standard setting that filters out templates with $\geq$ 90% sequence identity. This setup removes self-templates to prevent data leakage while retaining close structural homologs.
2. **Stringent Regime (ID $<$ 30%).** A stress test that filters out all templates with $\geq$ 30% sequence identity. This regime simulates "orphan" targets or de novo designs where no homologous structural information is available.

Template filtering was applied using a custom pre-processing script to generate separate input directories, which were then dynamically selected at runtime via the `--precomputed_templates_a3m_path` argument. In both regimes, a maximum of 4 top-ranked templates were passed to the model.

##### B.3 Statistical Analysis

To assess input feature quality, we measured MSA depth (raw count) and the effective number of sequences ($N_{\text{eff}}$). Structural prediction accuracy was quantified using TM-score and RMSD relative to ground truth. Model confidence was assessed using pLDDT and pTM.

Statistical comparisons were performed using the `SciPy` library [11]. Differences in metrics were initially evaluated using paired t-tests. To rigorously assess the functional equivalence of the two pipelines, we employed a **Two One-Sided Test (TOST)** [12] procedure with regime-specific criteria:

1. **Input Equivalence (Log-Ratio):** For MSA metrics ($N_{\text{eff}}$, Depth), which follow log-normal distributions, we performed a Log-Ratio TOST. We applied the standard bioequivalence margin of $[0.80, 1.25]$ [13], a rigorous threshold conventionally used to establish statistical equivalence in biological datasets.
2. **Output Equivalence (Absolute Difference):** For structural metrics (TM-score, RMSD, pLDDT), we applied *strict absolute margins* to ensure high fidelity. The equivalence bounds ($\epsilon$) were set based on community standards: $\epsilon = 0.02$ for TM-score and pTM, $\epsilon = 0.5$ Å for RMSD, and $\epsilon = 1.0$ for pLDDT.

A $p$-value $< 0.05$ in the TOST procedure indicates that the performance difference falls significantly within these specified equivalence bounds.

### Appendix C Supplementary Results

##### C.1 Template Retrieval Sensitivity and Structural Robustness

While the main text focuses on the functional equivalence of the final structures, we further analyzed the intermediate template retrieval steps and performance under a stringent evolutionary scenario (Figure C1).

**Template Retrieval Sensitivity.** We compared the ability of each pipeline to identify valid structural templates from the PDB70 database. Due to the heuristic nature of the accelerated search, we observed a trade-off in retrieval sensitivity. As shown in Figure C1a, the baseline hmmsearch pipeline retrieved templates for 27/32 targets, while the MMseqs2-GPU pipeline identified templates for 23/32 targets. Furthermore, Figure C1b indicates that for certain targets, MMseqs2-GPU retrieved templates with lower sequence identity compared to the exhaustive profile-based search of hmmsearch (points falling below the diagonal).

**Structural Robustness Under Stringent Filtering ($<$ 30%).** Crucially, however, this reduction in template quantity and quality did not propagate to the downstream prediction accuracy. To rigorously test this robustness, we evaluated performance under a "Stringent Regime" where all templates with $>$ 30% identity were excluded. As shown in Figure C1c-f, the global accuracy metrics remained strictly aligned despite the differences in template inputs. Quantitative analysis confirmed this equivalence: the mean difference in TM-score was negligible ($\Delta = -0.0015$), and RMSD variation was minimal ($\Delta = +0.0037$ Å), both falling well within strict equivalence bounds. This result strongly suggests that AF3 is robust to minor variations in template assignment and that AlphaFast provides sufficient evolutionary signal to maintain high-fidelity modeling even when template retrieval is suboptimal.

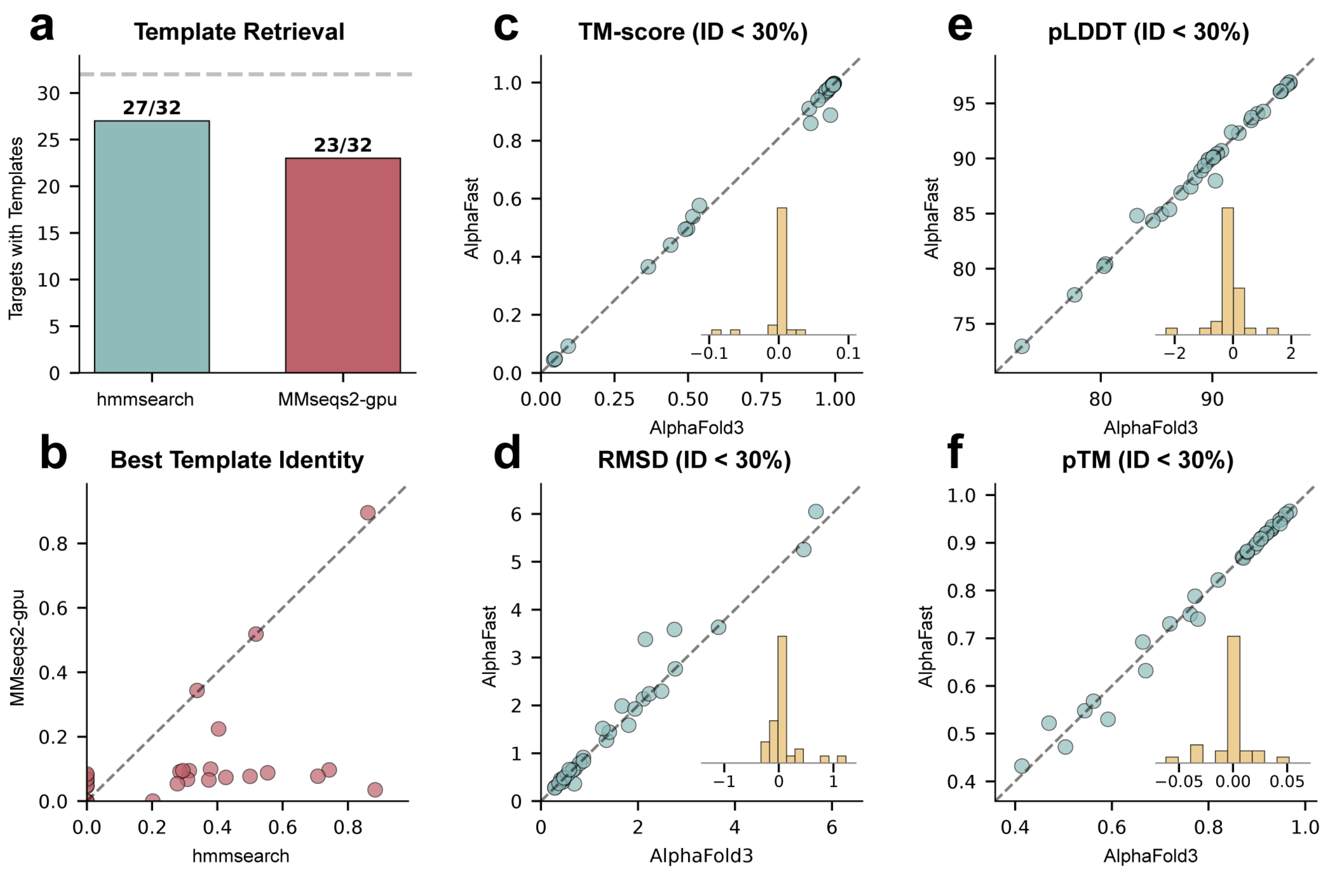


**Fig. C1 Template retrieval performance and structural accuracy under the Stringent Regime (< 30% Identity).** (a) Template retrieval success rate. MMseqs2-GPU shows slightly reduced sensitivity (23/32) compared to the exhaustive hmmsearch (27/32). (b) Comparison of the sequence identity of the best template found by each pipeline. Some targets show lower identity matches with MMseqs2-GPU (below diagonal). (c-f) Structural accuracy comparisons under the **Stringent Regime**. Remarkably, despite the reduction in template retrieval performance, MMseqs2-GPU maintains equivalent TM-score (c), RMSD (d), pLDDT (e), and pTM (f) to the baseline, demonstrating the robustness of the accelerated pipeline.

#### C.2 Statistical Bioequivalence Verification

To quantify the agreement between AlphaFast and the baseline JackHMMER pipeline, we performed a Two One-Sided Test (TOST) as detailed in Section B. Table C2 presents the comprehensive results of this assessment on the Accuracy Benchmark set ($N = 30$).

The analysis confirms that AlphaFast is statistically bioequivalent to the standard pipeline. For input features, while the heuristic search of MMseqs2 yields a Geometric Mean Ratio (GMR) of 87.1% for raw MSA depth, the effective sequence count ($N_{\text{eff}}$) is preserved with a GMR of 107.6%, indicating no loss of evolutionary signal. Consequently, all downstream structural metrics (TM-score, RMSD, pLDDT, and pTM) fall well within the strict equivalence margins, demonstrating that the speedup is achieved without compromising modeling accuracy.

**Table C2 Bioequivalence Assessment.** Input quantity was assessed using Geometric Mean Ratio (GMR) against standard bioequivalence limits (0.80–1.25). Structural fidelity was assessed via absolute mean difference ($\Delta$). Note that AlphaFast preserves effective information ($N_{\text{eff}}$) despite reducing raw depth.

| Metric | Comparison Metric | Equivalence Margin | Result |
|---|---|---|---|
| *Input Quantity (Log-Ratio TOST)* | | | |
| MSA Depth | GMR = 87.1% | $[0.80, 1.25]$ | Equivalent |
| $N_{\text{eff}}$ | GMR = 107.6% | $[0.80, 1.25]$ | Equivalent |
| *Output Quality (Absolute Difference TOST)* | | | |
| TM-score | $\Delta = +0.002$ | $\pm 0.02$ | Equivalent |
| RMSD (Å) | $\Delta = 0.00$ | $\pm 0.5$Å | Equivalent |
| pLDDT | $\Delta = -0.16$ | $\pm 2.0$ | Equivalent |
| pTM | $\Delta = -0.002$ | $\pm 0.02$ | Equivalent |

#### C.3 Serverless Inference Performance and Cost Efficiency

To ensure broad accessibility for researchers lacking dedicated HPC resources, we evaluated the cost-effectiveness of deploying AlphaFast on a serverless infrastructure (Modal). We benchmarked the performance of single-GPU instances—NVIDIA L40S versus NVIDIA H200—processing a cohort of 512 targets to simulate a production-scale high-throughput workload.

Table C3 summarizes the latency and cost metrics. When considering the GPU rental rate in isolation, the H200 incurs a slightly higher cost per input ($0.0357) compared to the L40S ($0.0349). However, because serverless providers bill for the concurrent usage of all provisioned resources (CPU and RAM), the H200's superior throughput—reducing average processing time from 64.3s to 28.3s ($\sim 2.3\times$ speedup)—leads to a significant reduction in auxiliary resource costs.

Consequently, the total cost per input (GPU + CPU + RAM) is lower on the H200 ($0.0385) than on the L40S ($0.0413). These results demonstrate that high-tier hardware like the H200 provides a dual advantage for serverless AlphaFast deployment, offering both drastic wall-clock time savings and superior overall cost efficiency for large-scale inference tasks.

We utilized Modal as the inference provider and calculated costs based on rates at the time of writing (February 2026).

**Table C3 Cost and Performance Comparison of Serverless Deployment (Modal).** Benchmarks were conducted using a cohort of 512 inputs. While the GPU-only cost is marginally higher for the H200, its 2.3× speedup reduces total system resource consumption, making it the more economical choice.

| Hardware | Time/Input (s) | GPU Cost /Input ($) | Total Cost /Input ($) | Total Cost (512 inputs) | Performance |
|---|---|---|---|---|---|
| NVIDIA L40S (Single) | 64.3 | 0.0349 | 0.0413 | $21.14 | Baseline |
| **NVIDIA H200 (Single)** | **28.3** | **0.0357** | **0.0385** | **$19.71** | 2.3× **Speedup** |

#### C.4 Code Availability

The source code for AlphaFast is openly available on GitHub at https://github.com/RomeroLab/alphafast.

#### C.5 Full Experimental Data

The benchmark datasets generated and analyzed during the current study are available in the Figshare repository, https://doi.org/10.6084/m9.figshare.31343287.

### References


[1] Suzek, B. E. *et al.* Uniref clusters: a comprehensive and scalable alternative for improving sequence similarity searches. *Bioinformatics* **31**, 926–932 (2015).

[2] Mitchell, A. L. *et al.* Mgnify: the microbiome analysis resource in 2020. *Nucleic acids research* **48**, D570–D578 (2020).

[3] Mirdita, M. *et al.* ColabFold: making protein folding accessible to all. *Nat. Methods* **19**, 679–682 (2022). URL https://www.nature.com/articles/s41592-022-01488-1.

[4] Jumper, J. *et al.* Highly accurate protein structure prediction with alphafold. *nature* **596**, 583–589 (2021).

[5] Consortium, U. Uniprot: a worldwide hub of protein knowledge. *Nucleic acids research* **47**, D506–D515 (2019).

[6] Eddy, S. R. Accelerated profile hmm searches. *PLoS computational biology* **7**, e1002195 (2011).

[7] Kallenborn, F. *et al.* GPU-accelerated homology search with MMseqs2. *Nat. Methods* **22**, 2024–2027 (2025). URL https://www.nature.com/articles/s41592-025-02819-8.

[8] wwPDB consortium. Protein data bank: the single global archive for 3d macromolecular structure data. *Nucleic Acids Research* **47**, D520–D528 (2018). URL https://doi.org/10.1093/nar/gky949.

[9] Bradbury, J. *et al.* Jax: composable transformations of python+ numpy programs (2018).

[10] Tillet, P., Kung, H.-T. & Cox, D. Triton: an intermediate language and compiler for tiled neural network computations (2019).

[11] Virtanen, P. *et al.* Scipy 1.0: fundamental algorithms for scientific computing in python. *Nature methods* **17**, 261–272 (2020).

[12] Koch, G. G. One-sided and two-sided tests and $\rho$ values. *Journal of biopharmaceutical statistics* **1**, 161–170 (1991).

[13] Schuirmann, D. J. A comparison of the two one-sided tests procedure and the power approach for assessing the equivalence of average bioavailability. *Journal of pharmacokinetics and biopharmaceutics* **15**, 657–680 (1987).
